# Investigating the roles for essential genes in the regulation of the circadian clock in *Synechococcus elongatus* using CRISPR interference

**DOI:** 10.1101/2022.10.10.511597

**Authors:** Nouneh Boodaghian, Hyunsook Park, Susan E. Cohen

## Abstract

Circadian rhythms are approximately 24-hour cycles that allow organisms to anticipate predictable environmental changes such as light intensity, temperature, and humidity. Circadian rhythms are found widely throughout nature where cyanobacteria are the simplest organisms known to carry out circadian rhythmicity. Circadian rhythmicity in cyanobacteria is carried out via the KaiA, KaiB and KaiC core oscillator proteins that keep ~24h time. A series of input and output proteins; CikA, SasA and RpaA, regulate the clock by sensing environmental changes, and time rhythmic activities, including global rhythms of gene expression. Our previous work identified a novel set of KaiC-interacting proteins, some of which are encoded by genes that are essential for viability. To understand the relationship of these essential genes to the clock we applied CRISPR interference (CRISPRi) which utilizes a deactivated Cas9 protein and single guide RNA (sgRNA) to reduce the expression of target genes but not fully abolish their expression to allow for survival. Eight candidate genes were targeted, and strains were analyzed by quantitative Reverse Transcriptase (RT)-PCR for reduction of gene expression and rhythms of gene expression were monitored to analyze circadian phenotypes. We present data that reduced expression of SynPCC942_0001, which encodes for the β-clamp of the replicative DNA polymerase, or SynPCC7942_1081, which likely encodes for KtrA involved in K^+^ transport, displayed a long circadian period compared to the wild type. As neither of these proteins have been previously implicated in the circadian clock, these data suggest that diverse cellular processes, DNA replication and K^+^ transport, can influence the circadian clock and represent new avenues to understand clock function.

## INTRODUCTION

Circadian rhythms are approximately 24-hour cycles that allow organisms to anticipate predictable environmental changes such as light intensity, temperature, and humidity. While the proteins that carry out circadian rhythmicity are different in each model system in which they are studied they all share the same fundamental principles of persistence, entrainment, and temperature compensation. Cyanobacteria are a group of photosynthetic bacteria and were the first organisms to perform oxygenic photosynthesis (Sanchez-Baracaldo & Cardona, 2020). Cyanobacteria are also known to be the simplest organisms known to possess a circadian clock, and *Synechococcus elongatus* PCC 7942 has emerged as the premier model system to elucidate the molecular details of the cyanobacterial circadian clock.

The cyanobacterial circadian clock is governed by three core oscillator proteins: KaiA, KaiB and KaiC. KaiC is a hexameric protein consisting of a CI and CII domains and functions as an autokinase, autophosphatase and ATPase (Nishiwaki et al., 2004; Xu et al., 2004). The CII domain of KaiC contains A-loops extensions that bind to KaiA during the day. The binding of KaiA to the A-loops at dawn promote KaiC autophosphorylation (Chang et al., 2012; Kim et al., 2008) on serine and threonine amino acids located in the CII domain in an ordered manner where threonine is phosphorylated first followed by serine (Nishiwaki et al., 2007; Nishiwaki et al., 2004; Rust et al., 2007). Once in the fully phosphorylated state, at dusk, KaiB will bind to the CI domain of KaiC and KaiA, sequestering KaiA away from the A-loops and promoting KaiC’s autophosphatase activity (Chang et al., 2012; Kim et al., 2008). This cycle of KaiC phosphorylation and dephosphorylation occurs over a ~24-h period and functions as the basic timekeeping mechanism in cyanobacteria. Deletion or over-expression of any of the *kai* genes abolishes the circadian clock (Ishiura et al., 1998).

The core oscillator receives input from the environment, through input proteins, and transmits temporal information, through output proteins, to clock controlled activities including, timing of cell division (Dong et al., 2010; Mori et al., 1996), compaction of the chromosome (Smith & Williams, 2006; Woelfle et al., 2007) and regulation of global patterns of gene expression (Ishiura et al., 1998; Kondo et al., 1994). CikA is a protein that plays key roles in both circadian input and output (Gutu & O’Shea, 2013; Schmitz et al., 2000). CikA monitors cellular redox through the binding of quinones in the membrane that allows the oscillator to synchronize with the environment, specifically signaling the night-time state (Ivleva et al., 2006; Kim et al., 2012). SasA, the histidine kinase, and RpaA, the cognate response regulator for SasA, are part of a two-component regulatory system that transmit signals from the oscillator to clock controlled outputs. SasA interacts with phosphorylated KaiC through its KaiB-like sensory domain (Iwasaki et al., 2000), which allows for SasA to autophosphorylate itself and transfer the phosphate group to RpaA (Takai et al., 2006). Phosphorylated RpaA, RpaA-P, is a DNA binding transcription factor and is also known as the master regulator of rhythmic gene expression (Markson et al., 2013). Without RpaA there would be no transmission of time from the KaiABC proteins to the clock outputs. CikA, functions as a phosphatase that removes the phosphate from RpaA, inactivating RpaA. SasA and CikA activity drive circadian oscillation in RpaA phosphorylation (Gutu & O’Shea, 2013).

In addition to changes in protein levels that occur over the course of the day (Kitayama et al., 2008), the clock undergoes an elegant orchestration in its subcellular localization patterns where KaiA and KaiC are found diffuse throughout the cell during the day and localized to a single pole of the cell at night, in a circadian fashion (Cohen et al., 2014). In order to determine the mechanism of KaiC localization, immunoprecipitation followed by mass-spectrometry was performed to identify proteins that interact specifically with KaiC in either a localized or delocalized state. Mutant variants of a YFP-KaiC fusion that displayed constitutive polar localization, KaiC-AA, mutation of the phosphorylated serine and threonine residues to alanine, or constitutive cytoplasmic localization, KaiC-AE, mutation of the phosphorylated serine and threonine residues to alanine and glutamic acid were used. 29 proteins were identified to interact with diffused KaiC or localized KaiC, but not both (McKnight, 2022). Fifteen of these proteins mapped to genes that were identified as being essential (Rubin et al., 2015) meaning that they are crucial for cell viability and replacement with an antibiotic resistance cassette was unsuccessful. Another two genes, SynPCC7942_1081 and SynPCC7942_0001, were identified as nonessential and sick respectively (Rubin et al., 2015); however, complete deletion via replacement with an antibiotic resistance cassette was not obtained, suggesting that these genes are indeed essential for viability. Here, we target eight of these genes for additional characterization using CRISPR interference (CRISPRi) to determine how depletion effects the circadian clock.

CRISPR encodes Cas9, an RNA-guided nuclease, and single guide RNA (sgRNA), consisting of CRISPR RNA (crRNA) and trans-activating crRNA (tracrRNA). Cas9 is then guided to the target genes by the single guide where the targeted DNA is cleaved (Ran et al., 2013). CRISPR interference (CRISPRi) was repurposed in 2013 for genome regulation rather than genome editing, by inactivating the catalytic function of Cas9 and using it for RNA-guided transcription regulation without genetically altering the targeted sequence (Qi et al., 2013). Unlike Cas9 used in CRISPR-Cas9 the deactivated Cas9, dCas9, protein does not pose nuclease activity. CRISPRi continues to form a partial fusion of crRNA and tracrRNA known as the single guide RNA (sgRNA) to mimic the natural duplex making up the dCas9-sgRNA complex. This complex binds to the desired sequence, such as the promoter, which blocks transcription initiation, or within the 5’untranslated region (UTR) or coding sequence, which blocks transcription elongation, resulting in the suppression of transcription. CRISPRi has been used to deplete expression of essential and nonessential genes in various organisms, including cyanobacteria (Gordon et al., 2016; Huang et al., 2016; Knoot et al., 2020).

Here we report that the decreased expression of two essential genes, SynPCC7942_0001, which encodes for the β-clamp needed for DNA replication, and SynPCC7942_1081, a hypothetical protein suggested to be part of the potassium transport system, display longer circadian rhythms of gene expression compared to the wildtype. Disruption to DNA replication or potassium transport had not been previously implicated in clock function, thus our results highlight how these previously unknown cellular processes may be connected to circadian clock function.

## MATERIALS AND METHODS

### Bacterial Strains and Growth Conditions

*Synechococcus elongatus* PCC 7942 obtained from the Golden lab at UCSD (Golden & Sherman, 1984) was grown in BG-11 media supplemented with appropriate antibiotics, at 30 °C under 50 to 300 μE of light for three to five days (Clerico et al., 2007). Strains were constructed by expressing genes in one of three *S. elongatus* neutral sites (NS) NS1, NS2 or NS3.

### CRISPR interference plasmid construction

Plasmids expressing dCas9 or sgRNAΦ, a plasmid expressing the sgRNA handle that binds to dCas9 without a target sequence, were obtained from Huang lab (Huang et al., 2016). dCas9 is expressed from NS1, under the P*_smtA_* promoter. dCas9 expression can be induced with Zinc Chloride (ZnCl_2_) from the P*_smtA_* promoter. sgRNAΦ is expressed from NS2 under the P_J23119_ constitutive promoter.

Primers to amplify the psgRNAΦ plasmid were designed to add a 20 bp target sequence, which targets either the 5’ UTR or within the open reading frame (ORF) of the genes of interest and are listed in Supplementary Table 1. For most genes, multiple target sequences were tested either within the 5’UTR, ORF or both. Q5 DNA polymerase (NEB Biolabs) was used for PCR amplification, followed by Dpn1 digest to remove the psgRNAΦ template lacking target sequence. Linear PCR fragments were then assembled into circular plasmids using GeneArt Seamless Cloning and Assembly Kit (Life Technologies) and propagated in *E. coli* XL1 Blue cells Plasmids were verified by Sanger sequencing. Plasmids generated are described in Supplementary Table 2.

### Circadian bioluminescence monitoring

Bioluminescence was monitored using a P*_kaiB_*-luciferase fusion reporter inserted into NS3 (AMC2158) (Cohen et al., 2018) under constant light and temperature (30° C). Cultures were entrained to 2-3 cycles of 12 hours light and 12 hours darkness at 30°C prior to synchronize the population as previously described (Mackey et al., 2007). In the cases where ZnCl_2_ was added, a final concentration of 8μM ZnCl_2_ was added after entrainment. Bioluminescence was monitored every 2 h from a TECAN Spark bioluminescence plate reader for 5-7 days. Data were plotted in excel and analyzed for rhythmicity using a MFourFit algorithm in BioDare2 (https://biodare2.ed.ac.uk)_(Zielinski et al., 2014).

### RNA Extraction

24-h prior to extraction ZnCl_2_ was added to a final concentration of 8 μM. 10 mL of culture at Optical Density (OD_750_) 0.2-0.4 was collected on ice and centrifuged at minus 10°C at 10,000 × g for 10 minutes. Pellets were resuspended with 1 mL of TriZol reagent and cells were lysed by10 rounds of vortexing for 30 second and incubating on ice for 30 seconds. RNA was isolated using a Direct-zol RNA Miniprep Kit (Zymo) per manufacturer’s instructions. DNase treatment was performed by adding 5μL DNase I along with 75 μL of DNA Digestion Buffer to the spin column and letting sit at room temperature for 15 mins. RNA concentration is determined via nanodrop and the quality of the RNA was determined by observing the 23S and 16S rRNA bands on 1.2 % agarose gel electrophoresis.

### Quantitative Real Time RT-PCR Analaysis

cDNAs were synthesized from 1 μg of total RNAs with Superscript IV Reverse Transcription Kit (Ambion) following the manufacturer’s protocol. Primers were designed using Primer3 (Koressaar et al., 2018; Koressaar & Remm, 2007; Untergasser et al., 2012) and listed in Supplemental Table 3. Quantitative real-time PCR was carried out using Maxima SYBR Green qPCR Master Mix (2X) (Molecular Biology) and Eppendorf Realplex System (Eppendorf) following the manufacturers’ protocol. *rpoA* gene was used as the endogenous control and the genomic DNA contamination was assessed by running no-RT control PCR with total RNA. The threshold cycle (Ct) values were obtained and the relative gene expression to the wild type strain was calculated by the 2^ΔΔCT^ method (Livak & Schmittgen, 2001) using the transcript level of *rpoA* as the endogenous control.

## RESULTS

### Construction of CRISPRi strains and evaluation by qRT-PCR

In order to establish CRISPRi we expressed dCas9 from NS1 under the P*_smtA_* inducible promoter and the sgRNA from NS2 under a constitutive promoter. Additionally, a luciferase reporter was expressed from NS3, which allowed us to monitor rhythms of gene expression (Figure 1). CRISPRi can be used to block either transcription initiation or transcription elongation, depending on where the sgRNA is designed to bind. In order to prevent possible off target effects, primers were designed to block transcription elongation. sgRNA plasmids containing 20 bp of target sequence homologous to where dCas9 was targeted, either within the 5’ UTR region or within the open reading frame (ORF) of the genes of interest were generated (Table 1). Of the 17 essential proteins identified as associating with KaiC, we identified 8 candidate genes to test for roles in the circadian clock using CRISPRi. Notably, we decided to not pursue ribosomal genes, as ribosomal proteins are typically highly abundant. Strains were generated to target both of these regions to maximize the chances of obtaining at least one strain with sufficient reduction in gene expression to be able to observe a phenotype, as differences in gene expression depending on where the sgRNA was targeted to have been observed (Huang et al., 2016). Two of the genes of interest were located within operons. For SynPCC7942_1081, the second gene of a two gene operon, and SynPCC7942_0507, is the second gene of a four gene operon, several targets within the ORF were selected so not to disrupt expression of the upstream genes. Although it is likely that gene expression was reduced for genes downstream of SynPCC7942_0507. We were able to successfully generate and propagate all CRISPRi strains, with the notable exception of 0001-5’UTR10, 1743-5’UTR3, 1743-5’UTR1, 2378-5’UTR22, which are sick and difficult to propagate even without inducing dCas9 with ZnCl_2_. As noted previously, there is leaky expression of dCas9 under non-inducing conditions and reduced gene expression was observed for candidate genes tested (Huang et al., 2016). This suggests that even mildly reduced expression of target genes in these strains’ effects viability.

**Figure 1.**
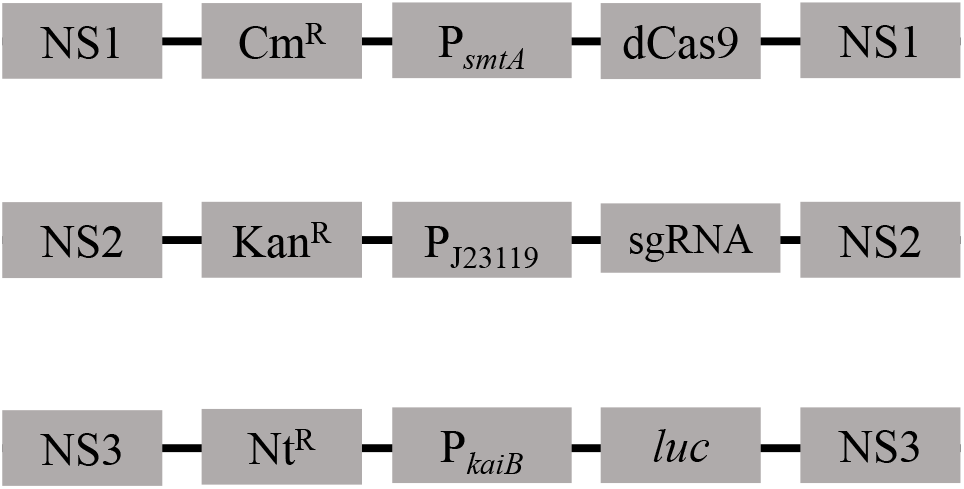
CRISPR interference Schematic. dCas9 is expressed in NS1 under the ZnCl_2_ inducible promoter P*_smtA_* with the chloramphenicol (Cm) antibiotic resistance cassette. The sgRNAs are expressed in NS2 under the constitutive promoter, P_J23119_, with kanamycin (Km) antibiotic resistance. The firefly luciferase reporter, P*_kaiB_-luc*, used to monitor circadian rhythms of gene expression is expressed from NS3 with the nourseothricin (Nt) antibiotic resistance cassette.

**Table 1.**
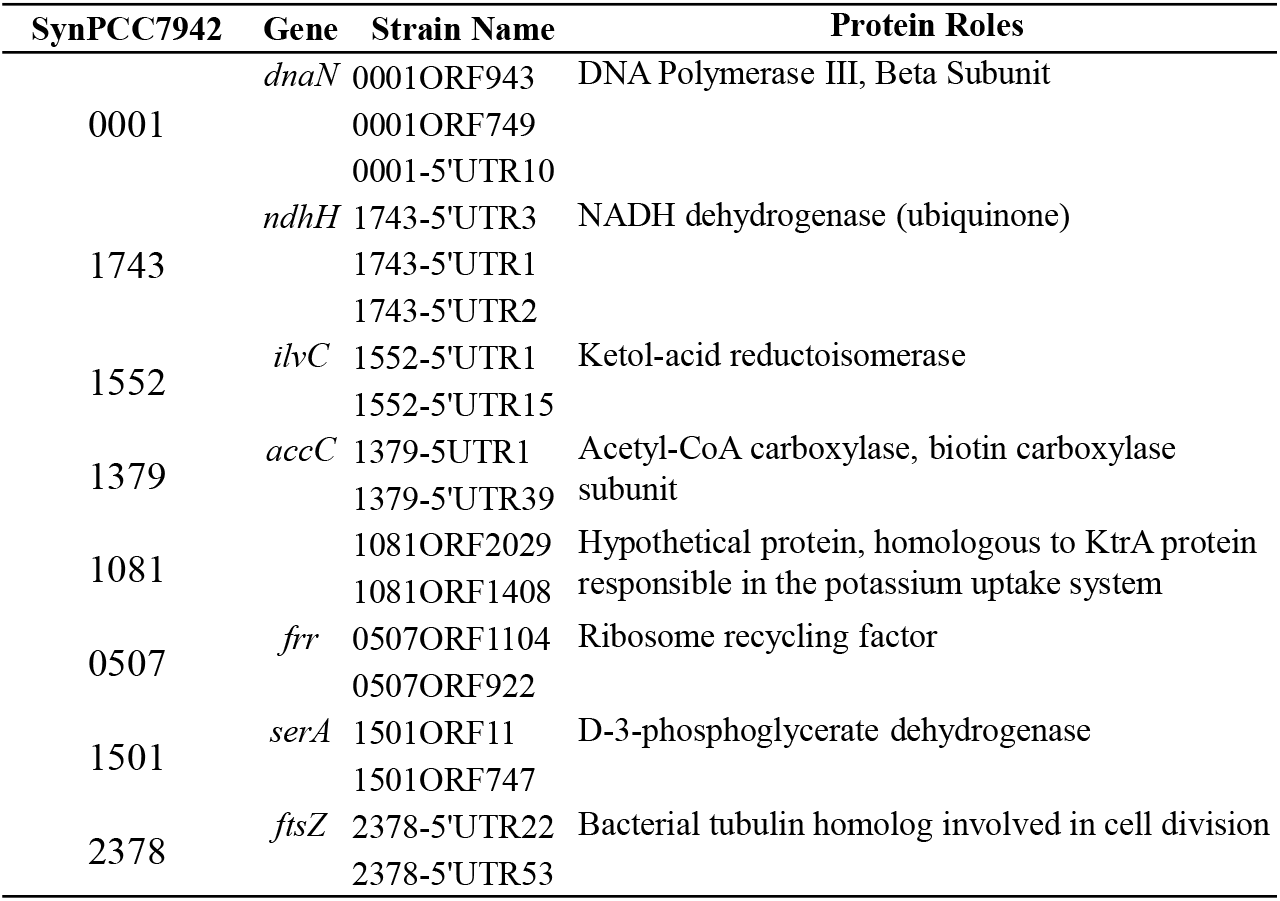
CRISPR interference Strain Constructs. sgRNA targets were constructed to bind to target sequences either within the 5’ UTR or the ORF of the gene of interest. The first column represents the SynPCC7942 gene number. The second column represents the gene name for the genes of interest, if available. The third column represents the CRISPRi strains constructed and can be broken down as such: the first four numbers are the SynPCC7942 gene number followed by the location to where the sgRNA guides the dCas9. The sgRNA targets were identified based on their position relative to the start codon of the gene of interest if the sgRNA targets the ORF, or to the transcription start site if the sgRNA targets within the 5’ UTR. The fourth column provides a description of the functionality of the proteins associated with the genes of interest.

To ensure that the CRISPRi resulted in reduced the gene expression of target genes, qRT-PCR was performed on all strains. Relative expression of the target genes in the CRISPRi strains compared to the wild type strain was calculated using *rpoA* as the endogenous control. We observed that the expression of target genes were significantly reduced in all CRISPRi strains under inducing conditions, although the magnitude of reduction varies depending on the sgRNA used (Figure 2).

**Figure 2.**
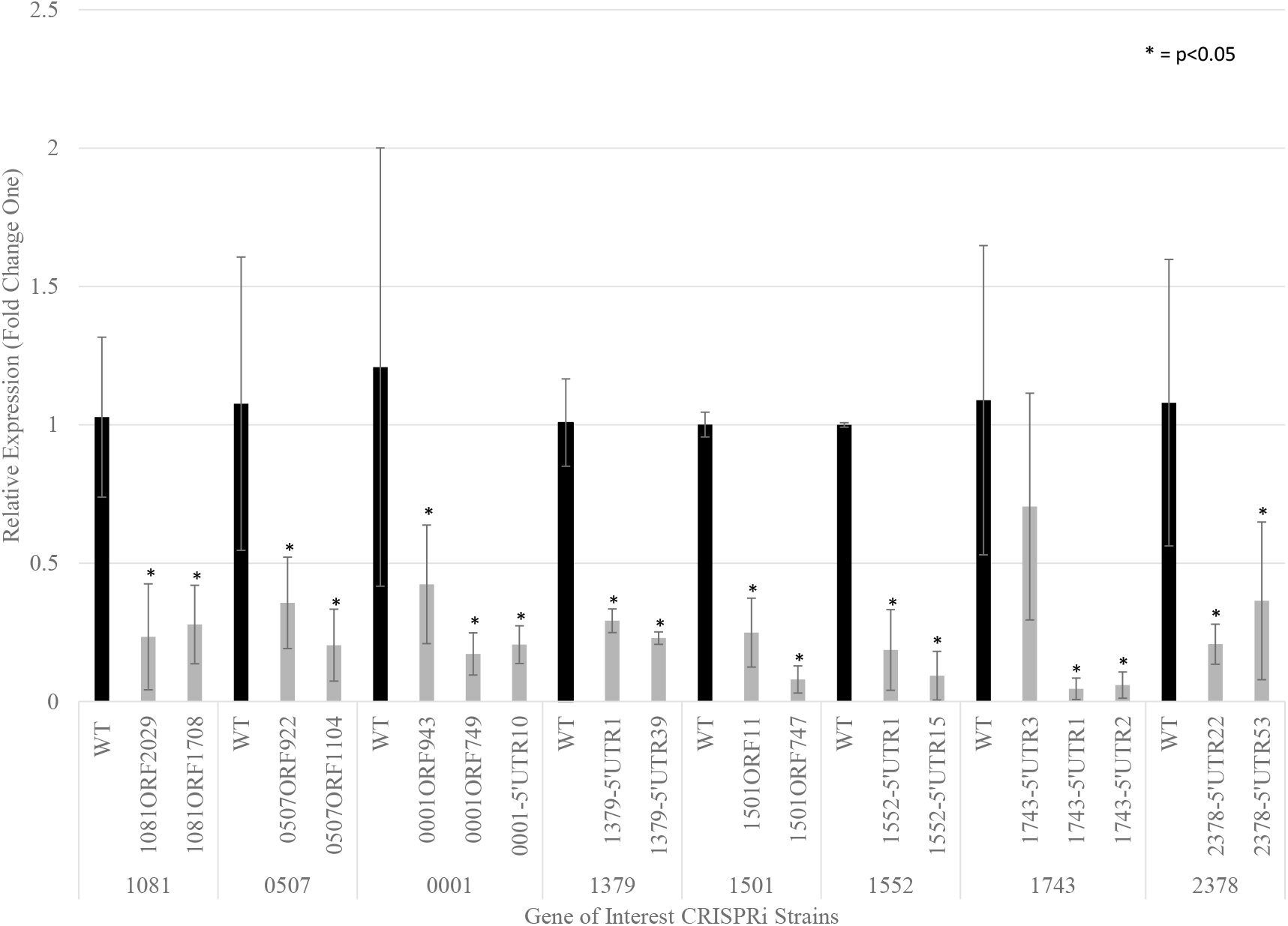
Confirmation of reduced gene expression by CRISPRi using quantitative PCR. *S. elongatus* genes expressed with CRISPR interference components (grey) compared to wild type expressing the corresponding genes (black). Results represent confirmation of reduced gene expression from CRISPRi strains. Standard deviations are marked by error bars, asterisk (*) indicated p-values of less than 0.05.

### Depletion of SynPCC7942_0001 and SynPCC7942_1081 results in altered circadian phenotypes

All strains were monitored for changes in circadian rhythms of gene expression using a luciferase reporter, under conditions were dCas9 was either induced with ZnCl_2_ or non-induced. We observe that the addition of 8 μM ZnCl_2_, required to induce dCas9 expression, did not affect circadian rhythms of gene expression (Figure 3). Of the 18 strains tested, 16 showed no phenotypic changes under non-induced conditions (Supplemental Table 4). The two strains that did show a change, both targeted SynPCC7942_0001, which resulted in long period rhythms compared to the wild type even without induction of dCas9 (Figure 4). In order to determine if induction of dCas9 could lead to enhanced circadian phenotypes, dCas9 was induced with ZnCl_2_. Indeed, more predominant phenotypic changes were observed when SynPCC7942_0001 was targeted (Figure 4). Additionally, a long circadian period was observed when SynPCC7942_1081 was targeted, whereas no obvious change was observed under non-inducing conditions (Figure 5). These data demonstrate that two of the candidate genes displayed phenotypic changes when targeted by the CRISPRi system, suggesting that reduced gene expression of these targets led to changes in the circadian clock.

**Figure 3.**
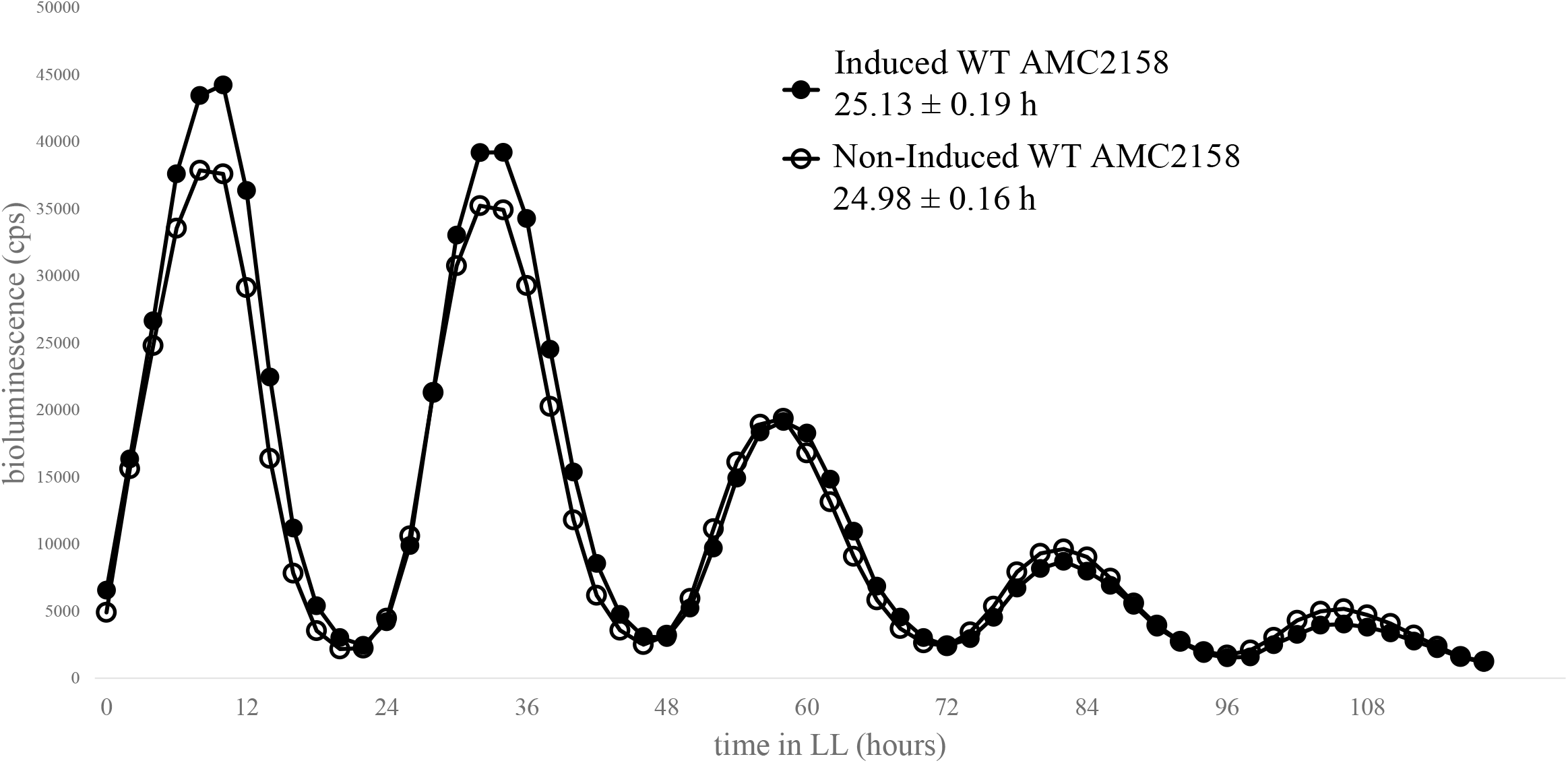
Induction of dCas9 with Zinc Chloride (ZnCl_2_) does not affect circadian period. Circadian rhythms of gene expression are monitored from a P_*kaiB*_-luciferase reporter either without dCas9 induction (open circle) or with dCas9 induction (closed circle) in the wild type strain (AMC2158). Period analysis shows that under induced conditions wild type period is 25.13 ± 0.19 h compared to non-induced conditions with a period of 24.98 ± 0.16 h, demonstrating that the addition of ZnCl_2_ does not affect circadian period.

**Figure 4.**
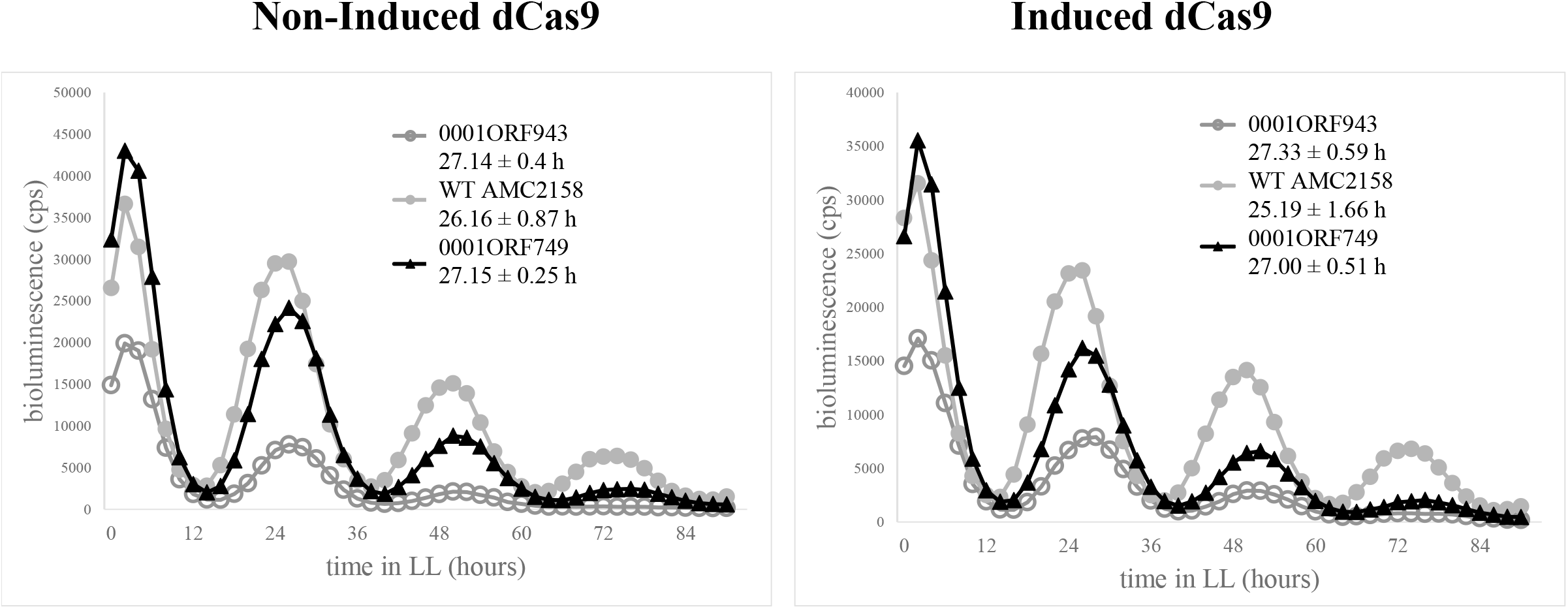
Targeting SynPCC7942_0001 with CRISPRi results in long circadian rhythms of gene expression. Circadian rhythms of gene expression monitored from a luciferase reporter either without dCas9 induction (Non-induced dCas9) or with dCas9 induction (Induced dCas9). CRISPRi strains (0001ORF943 & 0001ORF749) under non-induced conditions display a period of 27.14 ± 0.4 h (open, dark grey circle) and 27.15 ± 0.25 h (black, triangle) compared to the WT 26.16 ± 0.87 h (closed, light grey circle). CRISPRi strains (0001ORF943 & 0001ORF749) under induced conditions have a period of 27.00 ± 0.51 h (open, dark grey circle) and 27.33 ± 0.59 h (black triangle) compared to the WT 25.19 ± 1.66 h (closed, light grey circle). WT is AMC2158.

**Figure 5.**
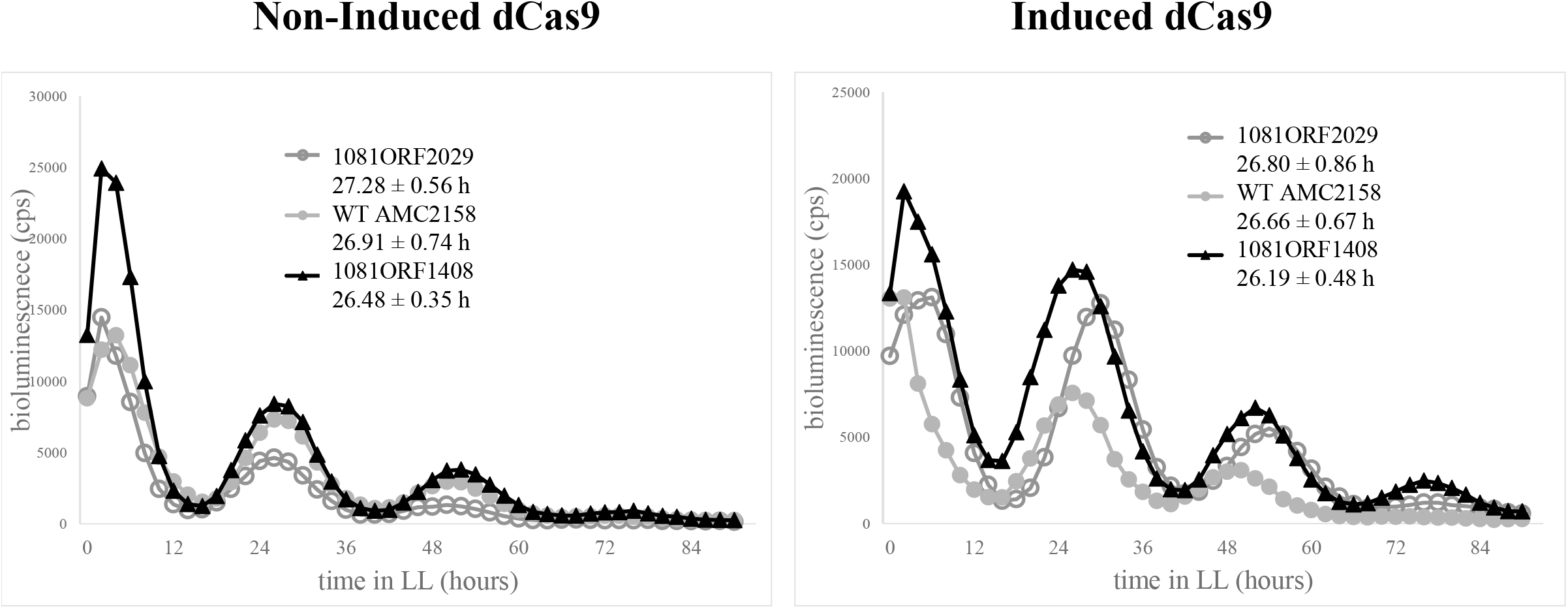
Circadian analysis of CRISPRi strains targeting SynPCC7942_1081. Circadian rhythms of gene expression monitored from a luciferase reporter either without dCas9 induction (Non-induced dCas9) or with dCas9 induction (Induced dCas9). CRISPRi strains (1081ORF2029 & 1081ORF1408) under noninduced conditions have a period of 27.28 ± 0.56 h (open, dark grey circle) and 26.48 ± 0.35 h (black triangle) compared to the WT 26.91 ± 0.74 h (closed, light grey circle). CRISPRi strains (1081ORF2029 & 1081ORF1408) under induced conditions have a period of 26.80 ± 0.86 h (open, dark grey circle) and 26.19 ± 0.48 h (black triangle) compared to the WT 26.66 ± 0.67 h (closed, light grey circle). WT is AMC2158.

Three CRISPRi strains were designed to target SynPCC7942_0001: 0001-5’UTR10, which targets within the 5’ UTR, and 0001ORF749 and 0001ORF943, both of which target within the open reading frame. As mentioned earlier, the strain 0001-5’UTR10 was sick and was difficult to propagate long enough to gather sufficient data. This suggests, that targeting the 5’UTR has a greater effect on gene expression compared to targets within the ORF, a trend we noticed for several genes. Period analysis shows that when SynPCC7942_0001 is targeted, a longer circadian period is observed for the two CRISPRi strains that target within the ORF. Without dCas9 induction we see period changes in both strains, strain 0001ORF749 has a period of 27.15 ± 0.25 h and strain 0001ORF943 has a period of 27.14 ± 0.4 h as compared to wild type which has a period of 26.16 ± 0.87 h (Figure 4). Upon dCas9 induction strain 0001ORF749 has a period of 27.00 ± 0.51 h and strain 0001ORF943 has a period of 27.33 ± 0.59 h compared to wild type which has a period of 25.19 ± 1.66 h demonstrating about a two-hour change in period (Figure 4).

Since SynPCC7942_1080 and SynPCC7942_1081 are part of an operon and shares the 5’ UTR we designed the CRISPRi strains targeting the ORF of SynPCC7942_1081 so not disrupt the expression of SynPCC7942_1080. Two CRISPRi strains targeting SynPCC7942_1081 were tested, 1081ORF2029 and 1081ORF1408. Without dCas9 induction, no major changes to the circadian clock were observed (Figure 5). However, upon dCas9 induction, long period rhythms were observed for both CRISPRi strains, although the effects were different depending on where the sgRNA was targeted. Strain 1081ORF2029 displayed a period of 26.80 ± 0.86 h and strain 1081ORF1408 displayed a period of 26.19 ± 0.49 h as compared to the wild type which has a period of 26.66 ± 0.67 h (Figure 5). These data demonstrate that targeting SynPCC7942_1081 results in changes to the circadian clock, where long period rhythms were observed. The strain that targeted a sequence closer to the 3’ end of the gene displayed greater phenotypes compared to sgRNA that targeted closer to the 5’ end of the gene.

### SynPCC7942_1081 likely encodes for a *ktrA* homolog

SynPCC7942_1081 is listed as hypothetical protein; however, genome annotation websites list peripheral membrane proteins, TrkA and KtrA, involved in K^+^ transport as possible homologs. TrkA is essential for the activity of the Trk potassium uptake system and has been described as mediating constitutive, low-affinity high-rate K^+^ transport energized by the protonmotive force (Rhoads & Epstein, 1977). Ktr is a sodium dependent potassium transport system which is crucial in playing a role in the response of hyperosmotic stress (Matsuda & Uozumi, 2006). Through protein alignments we observed that SynPCC7942_1081 had 17.56% identity to TrkA from *E. coli* and 29.57% identity to TrkA from *Synechococcus sp*. PCC 7002, a marine *Synechococcus* species, whereas *S. elongatus* PCC 7942 is a freshwater cyanobacterium (Figure 6). We then aligned SynPCC7942_1081 to the Ktr proteins KtrA, KtrB and KtrE from *Synechocystis sp*. PCC 6803. We observe that SynPCC7942_1081 has a 45.61% amino acid identity to the KtrA protein from *Synechocystis sp*. PCC 6803 (Figure 6). This led to the conclusion that SynPCC7942_1081 of *S. elongatus* most likely encodes for KtrA within the Ktr potassium transport system rather than a Trk potassium transport system. KtrA is proposed to regulate the potassium transport activity of KtrB, the membrane transporter, by changing its binding affinity from NAD^+^ to NADH through conformational change (Zulkifli et al., 2010).

**Figure 6.**
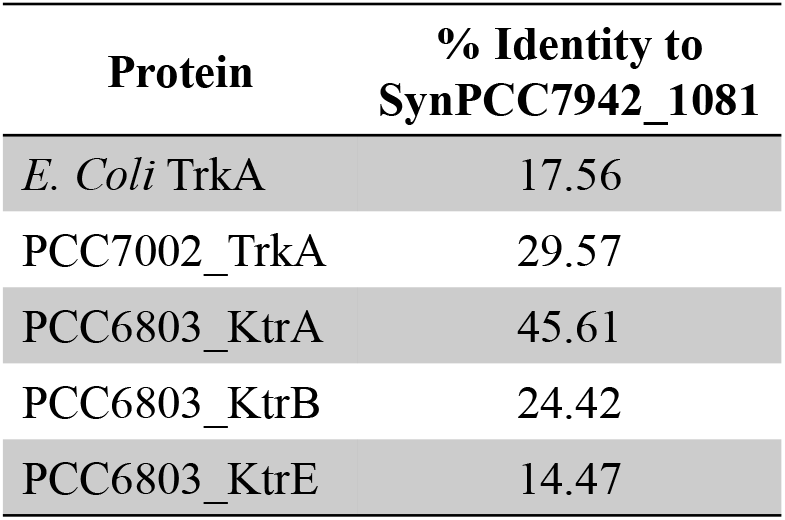
SynPCC7942_1081 shares homology with KtrA. Amino acid sequence of SynPCC7942_1081 was compared to TrkA from *E. coli* and *Synechococcus sp*. PCC 7002 along with KtrABE from *Synechocystis* PCC 6803. SynPCC7942_1081 has the highest percentage identity to the KtrA protein from *Synechocystis* PCC 6803 at 45.61%.

## Discussion

The utilization of CRISPRi here has allowed us to investigate the roles for essential genes in regulating the circadian clock of *S. elongatus*. Prior to CRISPRi, ways to deplete essential genes often resulted in inconsistent depletion. Here we targeted eight essential candidate genes, which had been identified as co-purifying with KaiC, and determined their effects on the cyanobacterial circadian clock. We were able to confirm through quantitative RT-PCR that reduction of gene expression was observed for all CRISPRi strains generated. We found that reduced expression of two genes, SynPCC7942_0001 and SynPCC7942_1081, resulted in longer circadian period phenotypes. Greater phenotypic changes were observed when SynPCC7942_1081 was targeted within the open reading frame and not within the 5’ UTR, suggesting that the CRISPRi complex could be more efficient at inhibiting elongating RNA Polymerase at these positions. For strains in which phenotypic changes were not observed either the genes targeted do not affect the circadian clock or that gene expression needs to be reduced even further for a phenotype to be observed. It is possible that targeting different regions within these genes could potentially lead to enhanced reduction of gene expression allowing for the observation of additional circadian phenotypes.

SynPCC7942_0001, *dnaN*, encodes for the β subunit, that makes up the β processivity clamp, of the replicative DNA polymerase and is required for DNA replication. We found that CRISPRi strains targeting SynPCC7942_0001 resulted in long-period circadian rhythms, under condition where dCas9 was either induced or non-induced. Interestingly, it has been shown that the circadian clock schedules rhythmic assembly of the replisome, and that there are high levels of initiation of DNA replication at dawn, in order to minimize incomplete replication forks at night (Liao & Rust, 2021). Our data suggest that in addition to the clock timing DNA replication that DNA replication also regulates the clock, as circadian phenotypes are observed when the expression of SynPCC7942_0001 is reduced. While there are many possible explanations for why changes to the circadian clock are observed when expression of *dnaN* is reduced, it is possible that KaiC, which is localized to the cell pole at night, sequesters the beta clamp at night as a mechanism to prevent DNA replication from occurring at night.

Through protein alignment we were able to provide evidence that SynPCC7942_1081, which was annotated as a hypothetical protein, likely encodes for KtrA which is part of the Ktr potassium transport system observed in other cyanobacteria like *Synechocystis* sp. KtrA is a membrane associated protein, that associates with the membrane bound K^+^ transporter KtrB. Interestingly, SynPCC7942_1081 was identified as interacting with KaiC in a localized state, suggesting KtrA/SynPCC7942_1081 could be part of the nighttime clock complex that forms a focus at or near the poles of cells (McKnight, 2022 In Preparation). The cyanobacterial circadian clock synchronizes with the environment indirectly by monitoring cellular redox. CikA and KaiA bind to oxidized quinones at night (Ivleva et al., 2006), signaling the onset of darkness (Ivleva et al., 2006; Kim et al., 2012; Wood et al., 2010). Whereas, KaiC binds to ATP, and can sense the changes in the ratio of ATP to ADP which gradually declines throughout the night, determine the length of night time period (Lin et al., 2014; Rust et al., 2011). As KtrA binds to NAD^+^/NADH (Zulkifli et al., 2010), it is possible KtrA association with the core oscillator represents another mechanism by which redox signals are conveyed to the clock.

In sum, we have successfully applied CRISPR interference as a tool to study the roles for essential genes in the circadian clock in *S. elongatus*. We generated several sgRNAs targeting eight essential genes, previously shown co-purify with KaiC, and were able to demonstrate that reduced expression of two genes resulted in circadian phenotypes. While the molecular mechanisms by which these factors regulate clock function are not investigated here, our work sheds light on previously unknown cellular factors that influence clock and paves the way for investigating the roles for other essential genes in circadian clock function.

## SUPPLEMENTAL LEGENDS

**Supplemental Table 1.**
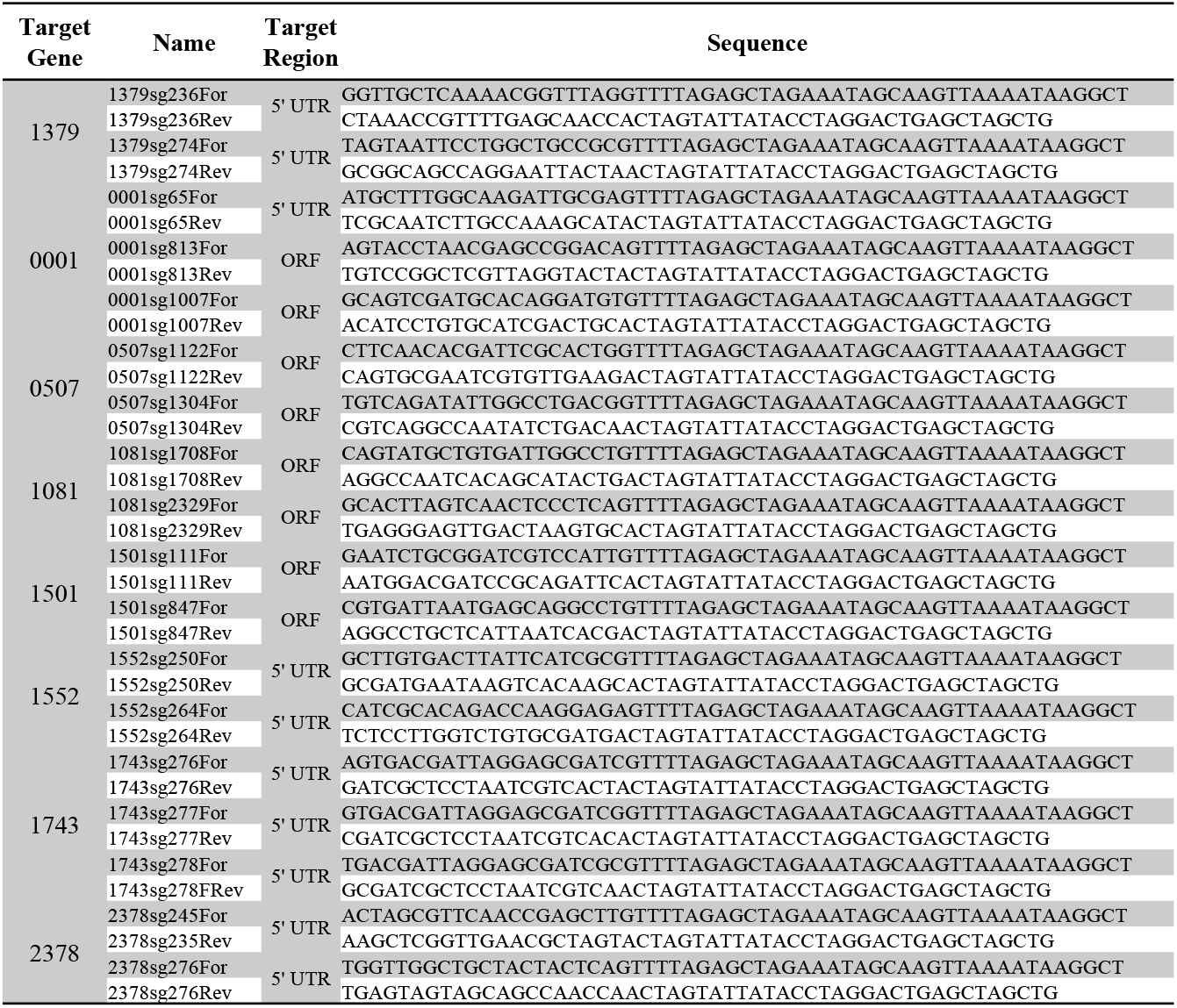
CRISPR interference single-guide RNA primer sequences. The first column represents the SynPCC number of the gene of interest. The second column represents primer names to amplify sgRNAΦ with the targeting sequence on 5’ and 3’ ends. The third column represents where within the gene was the sgRNA set to target and the fourth column is the full primer sequence.

**Supplemental Table 2.** CRISPR interference plasmid constructs. The first column represents the plasmid name. The second column represents the gene targeted by the sgRNA. The last column is the description of the plasmid including Neutral Site location, antibiotic resistance and sgRNA target location relative to the start of the ORF or 5’ UTR.

Boodaghian\_Supplementary Table 2
| Plasmid | Target Gene | Description |
| --- | --- | --- |
| pLA0029 | 1081 | NS2-Km-psgRNA::SynPCC7942 targeting within 1408 bases of the ORF |
| pLA0030 | 0001 | NS2-Km-psgRNA::SynPCC7942 targeting within 943 bases of the ORF |
| pLA0031 | 1501 | NS2-Km-psgRNA::SynPCC7942 targeting within 11 bases of the ORF |
| pLA0032 | 1743 | NS2-Km-psgRNA::SynPCC7942 targeting within 3 bases of the 5'UTR |
| pLA0033 | 0507 | NS2-Km-psgRNA::SynPCC7942 targeting within 1104 bases of the ORF |
| pLA0034 | 0507 | NS2-Km-psgRNA::SynPCC7942 targeting within 922 bases of the ORF |
| pLA0035 | 0001 | NS2-Km-psgRNA::SynPCC7942 targeting within 1 base of the 5'UTR |
| pLA0036 | 1379 | NS2-Km-psgRNA::SynPCC7942 targeting within 1 base of the 5'UTR |
| pLA0037 | 1379 | NS2-Km-psgRNA::SynPCC7942 targeting within 39 bases of the 5'UTR |
| pLA0038 | 1501 | NS2-Km-psgRNA::SynPCC7942 targeting within 747 bases of the ORF |
| pLA0039 | 1081 | NS2-Km-psgRNA::SynPCC7942 targeting within 2029 bases of the ORF |
| pLA0040 | 0001 | NS2-Km-psgRNA::SynPCC7942 targeting within 749 bases of the ORF |
| pLA0041 | 1552 | NS2-Km-psgRNA::SynPCC7942 targeting within 1 base of the 5'UTR |
| pLA0042 | 1743 | NS2-Km-psgRNA::SynPCC7942 targeting within 1 base of the 5'UTR |
| pLA0043 | 2378 | NS2-Km-psgRNA::SynPCC7942 targeting within 22 bases of the 5'UTR |
| pLA0044 | 1552 | NS2-Km-psgRNA::SynPCC7942 targeting within 15 bases of the 5'UTR |
| pLA0045 | 1743 | NS2-Km-psgRNA::SynPCC7942 targeting within 2 bases of the 5'UTR |
| pLA0046 | 2378 | NS2-Km-psgRNA::SynPCC7942 targeting within 1 base of the 5'UTR |

**Supplemental Table 3.**
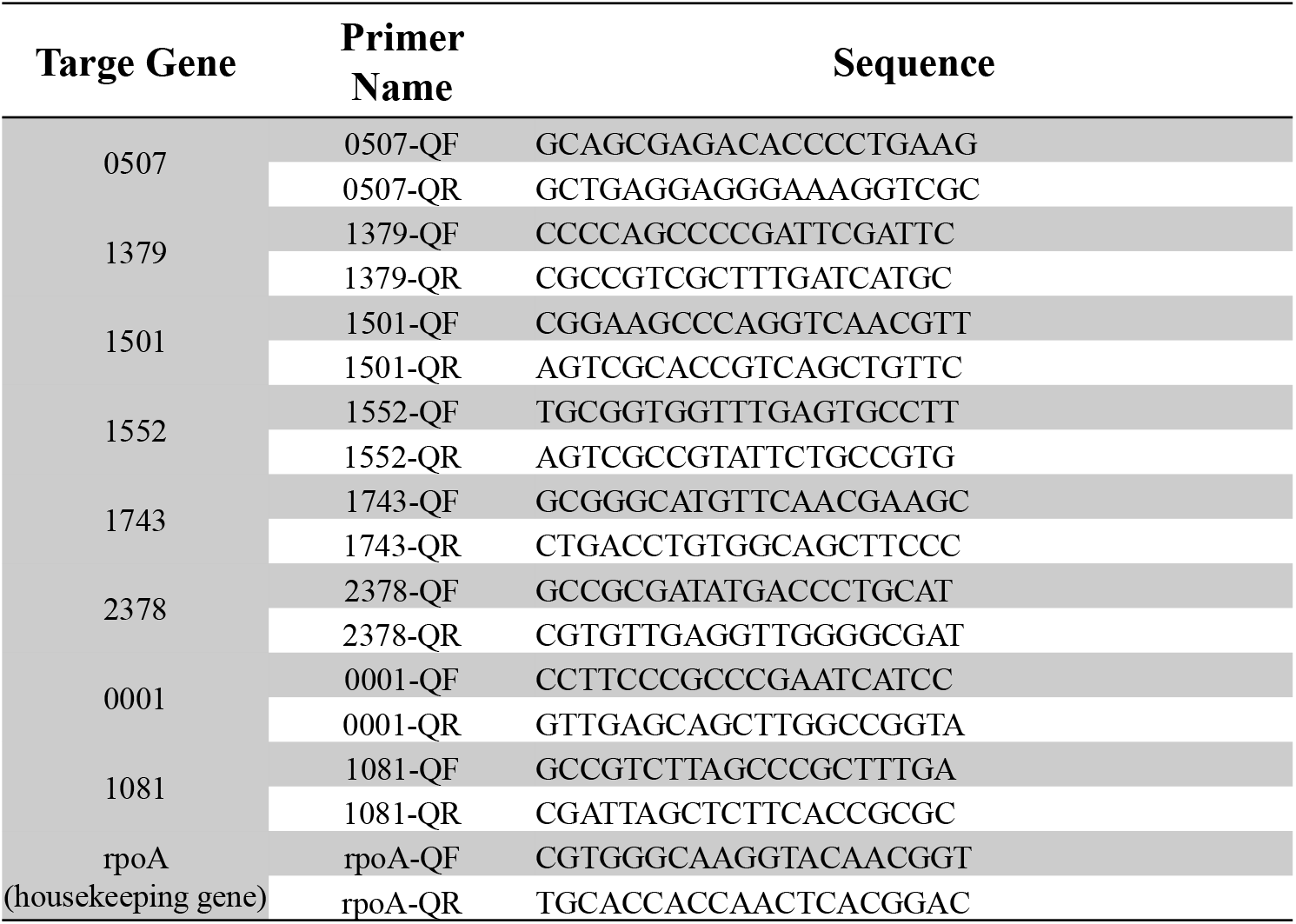
Quantitative PCR primer sequences. The first column represents the SynPCC7942 number of our target genes. The second column is the primer name assigned at the time of creation and lastly the fourth column is the full primer sequence.

**Supplemental Table 4.** Period analysis of CRISPR interference strains that did not display circadian phenotypes. The columns labeled “Strain” represent the strains for which period analysis is provided. The columns labeled “Time in LL (hours)” are the circadian periods calculated using BioDare2 represented in hours in constant light, under induced conditions.

Boodaghian\_Supplementary Table 4
| Strain | Time in LL (hours) | Strain | Time in LL (hours) |
| --- | --- | --- | --- |
| WT AMC2158 | 25.41 | 1501ORF747 | 25.82 |
| 1081ORF2029 | 25.55 | 1552-5'UTR1 | 25.64 |
| 1081ORF1408 | 25.65 | 1552-5'UTR15 | 26.09 |
| 0507ORF922 | 25.76 | 1743-5'UTR3 | 25.64 |
| 0507ORF1104 | 25.60 | 1743-5'UTR1 | 25.26 |
| 0001-5'UTR10 | 25.05 | 2378-5'UTR22 | 25.69 |
| 1379-5'UTR1 | 25.70 | 1743-5'UTR2 | 25.92 |
| 1379-5'UTR39 | 25.95 | 2378-5'UTR53 | 25.59 |
| 1501ORF11 | 26.35 |  |  |

## REFERENCES

Chang, Y. G., Tseng, R., Kuo, N. W., & LiWang, A. (2012). Rhythmic ring-ring stacking drives the circadian oscillator clockwise. Proc Natl Acad Sci U S A, 109(42), 16847–16851. https://doi.org/10.1073/pnas.1211508109

Clerico, E. M., Ditty, J. L., & Golden, S. S. (2007). Specialized techniques for site-directed mutagenesis in cyanobacteria. Methods Mol Biol, 362, 155–171. https://doi.org/10.1007/978-1-59745-257-1_11

Cohen, S. E., Erb, M. L., Selimkhanov, J., Dong, G., Hasty, J., Pogliano, J., & Golden, S. S. (2014). Dynamic localization of the cyanobacterial circadian clock proteins. Curr Biol, 24(16), 1836–1844. https://doi.org/10.1016/j.cub.2014.07.036

Cohen, S. E., McKnight, B. M., & Golden, S. S. (2018). Roles for ClpXP in regulating the circadian clock in Synechococcus elongatus. Proc Natl Acad Sci U S A, 115(33), E7805–E7813. https://doi.org/10.1073/pnas.1800828115

Dong, G., Yang, Q., Wang, Q., Kim, Y. I., Wood, T. L., Osteryoung, K. W., van Oudenaarden, A., & Golden, S. S. (2010). Elevated ATPase activity of KaiC applies a circadian checkpoint on cell division in Synechococcus elongatus. Cell, 140(4), 529–539. https://doi.org/10.1016/j.cell.2009.12.042

Golden, S. S., & Sherman, L. A. (1984). Optimal conditions for genetic transformation of the cyanobacterium Anacystis nidulans R2. J Bacteriol, 158(1), 36–42. https://doi.org/10.1128/jb.158.1.36-42.1984

Gordon, G. C., Korosh, T. C., Cameron, J. C., Markley, A. L., Begemann, M. B., & Pfleger, B. F. (2016). CRISPR interference as a titratable, trans-acting regulatory tool for metabolic engineering in the cyanobacterium Synechococcus sp. strain PCC 7002. Metab Eng, 38, 170–179. https://doi.org/10.1016/j.ymben.2016.07.007

Gutu, A., & O’Shea, E. K. (2013). Two antagonistic clock-regulated histidine kinases time the activation of circadian gene expression. Mol Cell, 50(2), 288–294. https://doi.org/10.1016/j.molcel.2013.02.022

Huang, C. H., Shen, C. R., Li, H., Sung, L. Y., Wu, M. Y., & Hu, Y. C. (2016). CRISPR interference (CRISPRi) for gene regulation and succinate production in cyanobacterium S. elongatus PCC 7942. Microb Cell Fact, 15(1), 196. https://doi.org/10.1186/s12934-016-0595-3

Ishiura, M., Kutsuna, S., Aoki, S., Iwasaki, H., Andersson, C. R., Tanabe, A., Golden, S. S., Johnson, C. H., & Kondo, T. (1998). Expression of a gene cluster kaiABC as a circadian feedback process in cyanobacteria. Science, 281(5382), 1519–1523. https://doi.org/10.1126/science.281.5382.1519

Ivleva, N. B., Gao, T., LiWang, A. C., & Golden, S. S. (2006). Quinone sensing by the circadian input kinase of the cyanobacterial circadian clock. Proc Natl Acad Sci U S A, 103(46), 17468–17473. https://doi.org/10.1073/pnas.0606639103

Iwasaki, H., Williams, S. B., Kitayama, Y., Ishiura, M., Golden, S. S., & Kondo, T. (2000). A kaiC-interacting sensory histidine kinase, SasA, necessary to sustain robust circadian oscillation in cyanobacteria. Cell, 101(2), 223–233. https://doi.org/10.1016/S0092-8674(00)80832-6

Kim, Y. I., Dong, G., Carruthers, C. W., Jr., Golden, S. S., & LiWang, A. (2008). The day/night switch in KaiC, a central oscillator component of the circadian clock of cyanobacteria. Proc Natl Acad Sci U S A, 105(35), 12825–12830. https://doi.org/10.1073/pnas.0800526105

Kim, Y. I., Vinyard, D. J., Ananyev, G. M., Dismukes, G. C., & Golden, S. S. (2012). Oxidized quinones signal onset of darkness directly to the cyanobacterial circadian oscillator. Proc Natl Acad Sci U S A, 109(44), 17765–17769. https://doi.org/10.1073/pnas.1216401109

Kitayama, Y., Nishiwaki, T., Terauchi, K., & Kondo, T. (2008). Dual KaiC-based oscillations constitute the circadian system of cyanobacteria. Genes Dev, 22(11), 1513–1521. https://doi.org/10.1101/gad.1661808

Knoot, C. J., Biswas, S., & Pakrasi, H. B. (2020). Tunable Repression of Key Photosynthetic Processes Using Cas12a CRISPR Interference in the Fast-Growing Cyanobacterium Synechococcus sp. UTEX 2973. ACS Synth Biol, 9(1), 132–143. https://doi.org/10.1021/acssynbio.9b00417

Kondo, T., Tsinoremas, N. F., Golden, S. S., Johnson, C. H., Kutsuna, S., & Ishiura, M. (1994). Circadian clock mutants of cyanobacteria. Science, 266(5188), 1233–1236. https://doi.org/10.1126/science.7973706

Koressaar, T., Lepamets, M., Kaplinski, L., Raime, K., Andreson, R., & Remm, M. (2018). Primer3_masker: integrating masking of template sequence with primer design software. Bioinformatics, 34(11), 1937–1938. https://doi.org/10.1093/bioinformatics/bty036

Koressaar, T., & Remm, M. (2007). Enhancements and modifications of primer design program Primer3. Bioinformatics, 23(10), 1289–1291. https://doi.org/10.1093/bioinformatics/btm091

Liao, Y., & Rust, M. J. (2021). The circadian clock ensures successful DNA replication in cyanobacteria. Proc Natl Acad Sci U S A, 118(20). https://doi.org/10.1073/pnas.2022516118

Lin, J., Chew, J., Chockanathan, U., & Rust, M. J. (2014). Mixtures of opposing phosphorylations within hexamers precisely time feedback in the cyanobacterial circadian clock. Proc Natl Acad Sci U S A, 111(37), E3937–3945. https://doi.org/10.1073/pnas.1408692111

Livak, K. J., & Schmittgen, T. D. (2001). Analysis of relative gene expression data using realtime quantitative PCR and the 2(-Delta Delta C(T)) Method. Methods, 25(4), 402–408. https://doi.org/10.1006/meth.2001.1262

Mackey, S. R., Ditty, J. L., Clerico, E. M., & Golden, S. S. (2007). Detection of rhythmic bioluminescence from luciferase reporters in cyanobacteria. Methods Mol Biol, 362, 115–129. https://doi.org/10.1007/978-1-59745-257-1_8

Markson, J. S., Piechura, J. R., Puszynska, A. M., & O’Shea, E. K. (2013). Circadian control of global gene expression by the cyanobacterial master regulator RpaA. Cell, 155(6), 1396–1408. https://doi.org/10.1016/j.cell.2013.11.005

Matsuda, N., & Uozumi, N. (2006). Ktr-mediated potassium transport, a major pathway for potassium uptake, is coupled to a proton gradient across the membrane in Synechocystis sp. PCC 6803. Biosci Biotechnol Biochem, 70(1), 273–275. https://doi.org/10.1271/bbb.70.273

McKnight, B. M. K., S.; Le, T.H.; Carbonel, G.; Rodriguez, E.; Tran, A.L.; Duncan, N.R.; Golden, S.S.; Cohen, S.E. (2022). Roles for the RNA binding protein, Rbp2, in regulating the circadian clock in Synechococcus elongatus. In.

Mori, T., Binder, B., & Johnson, C. H. (1996). Circadian gating of cell division in cyanobacteria growing with average doubling times of less than 24 hours. Proc Natl Acad Sci U S A, 93(19), 10183–10188. https://doi.org/10.1073/pnas.93.19.10183

Nishiwaki, T., Satomi, Y., Kitayama, Y., Terauchi, K., Kiyohara, R., Takao, T., & Kondo, T. (2007). A sequential program of dual phosphorylation of KaiC as a basis for circadian rhythm in cyanobacteria. EMBO J, 26(17), 4029–4037. https://doi.org/10.1038/sj.emboj.7601832

Nishiwaki, T., Satomi, Y., Nakajima, M., Lee, C., Kiyohara, R., Kageyama, H., Kitayama, Y., Temamoto, M., Yamaguchi, A., Hijikata, A., Go, M., Iwasaki, H., Takao, T., & Kondo, T. (2004). Role of KaiC phosphorylation in the circadian clock system of Synechococcus elongatus PCC 7942. Proc Natl Acad Sci U S A, 101(38), 13927–13932. https://doi.org/10.1073/pnas.0403906101

Qi, L. S., Larson, M. H., Gilbert, L. A., Doudna, J. A., Weissman, J. S., Arkin, A. P., & Lim, W. A. (2013). Repurposing CRISPR as an RNA-guided platform for sequence-specific control of gene expression. Cell, 152(5), 1173–1183. https://doi.org/10.1016/j.cell.2013.02.022

Ran, F. A., Hsu, P. D., Wright, J., Agarwala, V., Scott, D. A., & Zhang, F. (2013). Genome engineering using the CRISPR-Cas9 system. Nat Protoc, 8(11), 2281–2308. https://doi.org/10.1038/nprot.2013.143

Rhoads, D. B., & Epstein, W. (1977). Energy coupling to net K+ transport in Escherichia coli K-12. J Biol Chem, 252(4), 1394–1401. https://www.ncbi.nlm.nih.gov/pubmed/320207

Rubin, B. E., Wetmore, K. M., Price, M. N., Diamond, S., Shultzaberger, R. K., Lowe, L. C., Curtin, G., Arkin, A. P., Deutschbauer, A., & Golden, S. S. (2015). The essential gene set of a photosynthetic organism. Proc Natl Acad Sci U S A, 112(48), E6634–6643. https://doi.org/10.1073/pnas.1519220112

Rust, M. J., Golden, S. S., & O’Shea, E. K. (2011). Light-driven changes in energy metabolism directly entrain the cyanobacterial circadian oscillator. Science, 331(6014), 220–223. https://doi.org/10.1126/science.1197243

Rust, M. J., Markson, J. S., Lane, W. S., Fisher, D. S., & O’Shea, E. K. (2007). Ordered phosphorylation governs oscillation of a three-protein circadian clock. Science, 318(5851), 809–812. https://doi.org/10.1126/science.1148596

Sanchez-Baracaldo, P., & Cardona, T. (2020). On the origin of oxygenic photosynthesis and Cyanobacteria. New Phytol, 225(4), 1440–1446. https://doi.org/10.1111/nph.16249

Schmitz, O., Katayama, M., Williams, S. B., Kondo, T., & Golden, S. S. (2000). CikA, a bacteriophytochrome that resets the cyanobacterial circadian clock. Science, 289(5480), 765–768. https://doi.org/10.1126/science.289.5480.765

Smith, R. M., & Williams, S. B. (2006). Circadian rhythms in gene transcription imparted by chromosome compaction in the cyanobacterium Synechococcus elongatus. Proc Natl Acad Sci U S A, 103(22), 8564–8569. https://doi.org/10.1073/pnas.0508696103

Takai, N., Nakajima, M., Oyama, T., Kito, R., Sugita, C., Sugita, M., Kondo, T., & Iwasaki, H. (2006). A KaiC-associating SasA-RpaA two-component regulatory system as a major circadian timing mediator in cyanobacteria. Proc Natl Acad Sci U S A, 103(32), 12109–12114. https://doi.org/10.1073/pnas.0602955103

Untergasser, A., Cutcutache, I., Koressaar, T., Ye, J., Faircloth, B. C., Remm, M., & Rozen, S. G. (2012). Primer3--new capabilities and interfaces. Nucleic Acids Res, 40(15), e115. https://doi.org/10.1093/nar/gks596

Woelfle, M. A., Xu, Y., Qin, X., & Johnson, C. H. (2007). Circadian rhythms of superhelical status of DNA in cyanobacteria. Proc Natl Acad Sci U S A, 104(47), 18819–18824. https://doi.org/10.1073/pnas.0706069104

Wood, T. L., Bridwell-Rabb, J., Kim, Y. I., Gao, T., Chang, Y. G., LiWang, A., Barondeau, D. P., & Golden, S. S. (2010). The KaiA protein of the cyanobacterial circadian oscillator is modulated by a redox-active cofactor. Proc Natl Acad Sci U S A, 107(13), 5804–5809. https://doi.org/10.1073/pnas.0910141107

Xu, Y., Mori, T., Pattanayek, R., Pattanayek, S., Egli, M., & Johnson, C. H. (2004). Identification of key phosphorylation sites in the circadian clock protein KaiC by crystallographic and mutagenetic analyses. Proc Natl Acad Sci U S A, 101(38), 13933–13938. https://doi.org/10.1073/pnas.0404768101

Zielinski, T., Moore, A. M., Troup, E., Halliday, K. J., & Millar, A. J. (2014). Strengths and limitations of period estimation methods for circadian data. PLoS One, 9(5), e96462. https://doi.org/10.1371/journal.pone.0096462

Zulkifli, L., Akai, M., Yoshikawa, A., Shimojima, M., Ohta, H., Guy, H. R., & Uozumi, N. (2010). The KtrA and KtrE subunits are required for Na+-dependent K+ uptake by KtrB across the plasma membrane in Synechocystis sp. strain PCC 6803. J Bacteriol, 192(19), 5063–5070. https://doi.org/10.1128/JB.00569-10

